# Orofacial behaviors, not eye movements, govern neural activity in mouse visual cortex

**DOI:** 10.64898/2026.02.04.703800

**Authors:** Atika Syeda, Miguel Angel Nunez-Ochoa, Lin Zhong, Marius Pachitariu, Carsen Stringer

## Abstract

Neural activity in mouse primary visual cortex (V1) correlates strongly with orofacial movements. Such strong modulation has not been found in the primate visual cortex during eye fixation [1], which led to the suggestion that the modulation may primarily depend on eye movements in both species [2]. Here we examined the influence of eye movements on neural activity in mouse visual cortex both in complete darkness and in the presence of different types of visual input. In all cases, we found that eye movements explain a smaller fraction of neural activity variance compared to orofacial behaviors. Additionally, we found that eye movements were correlated to orofacial movements, such as whisking and sniffing, and thus may be indirectly correlated to neural activity. These results further emphasize the impact of movement signals on mouse visual cortex during free viewing behavior.

---

Locomotion and other movements modulate neural activity during instructed and uninstructed behaviors in the sensory brain regions of flies [3, 4], zebrafish [5] and mice [6–9]. In mouse V1, a large fraction of the spontaneous neural activity is correlated to various orofacial movements, such as whisking and sniffing [7, 9], and locomotion leads to increased neural responses to visual stimuli [6]. In contrast, locomotion may have a slightly suppressive effect in primate V1 (but see [11]). Furthermore, representations related to spontaneous body or orofacial movements have not been found in primate V1 [1]. These apparently contrasting findings suggest differences in sensorimotor integration across species. However, a recent review raises the possibility that differences could be related to other signals, such as eye movements [2], which are typically unaccounted for by studies in mice [7, 9], but tightly controlled in non-human primate studies. Yet another possibility is that tight control of eye movements reduces orofacial behaviors overall, which may in turn reduce or eliminate their impact on neural activity.

Although eye movement-related neural signals have been well-studied in primates [1, 12, 13], they remain to be further characterized in rodents. In addition, primate visual cortical regions receive movement corollary discharge during fast saccadic eye movements [14]. In mice, different types of eye movements are observed such as saccades and vestibulo-ocular reflexes, which have homologs in primates [15]. In addition, eye movements drive activity in V1 [16–18] and in higher-order visual areas [19, 20]. However, the relative effect of eye movements and orofacial movements on neural activity in mouse visual cortex remains to be studied.

To investigate the contribution of eye and orofacial movements to neural activity, we presented images on three screens while mice were head-fixed on a cylindrical treadmill to allow for locomotion (Figure 1a). We recorded the activity of tens of thousands of visual cortical neurons using two-photon mesososcopic calcium imaging (Figure 1b, [21]), and we simultaneously monitored orofacial behaviors and eye movements with a camera (Figure 1c). To analyze the behavior, we extracted principal components (PCs) of behavioral videos and tracked the pupil position by computing its center-of-mass [9]. These behavioral readouts formed the inputs to deep convolutional neural networks which we trained to predict neural activity (Figure 1d) [9].

**Figure 1:**
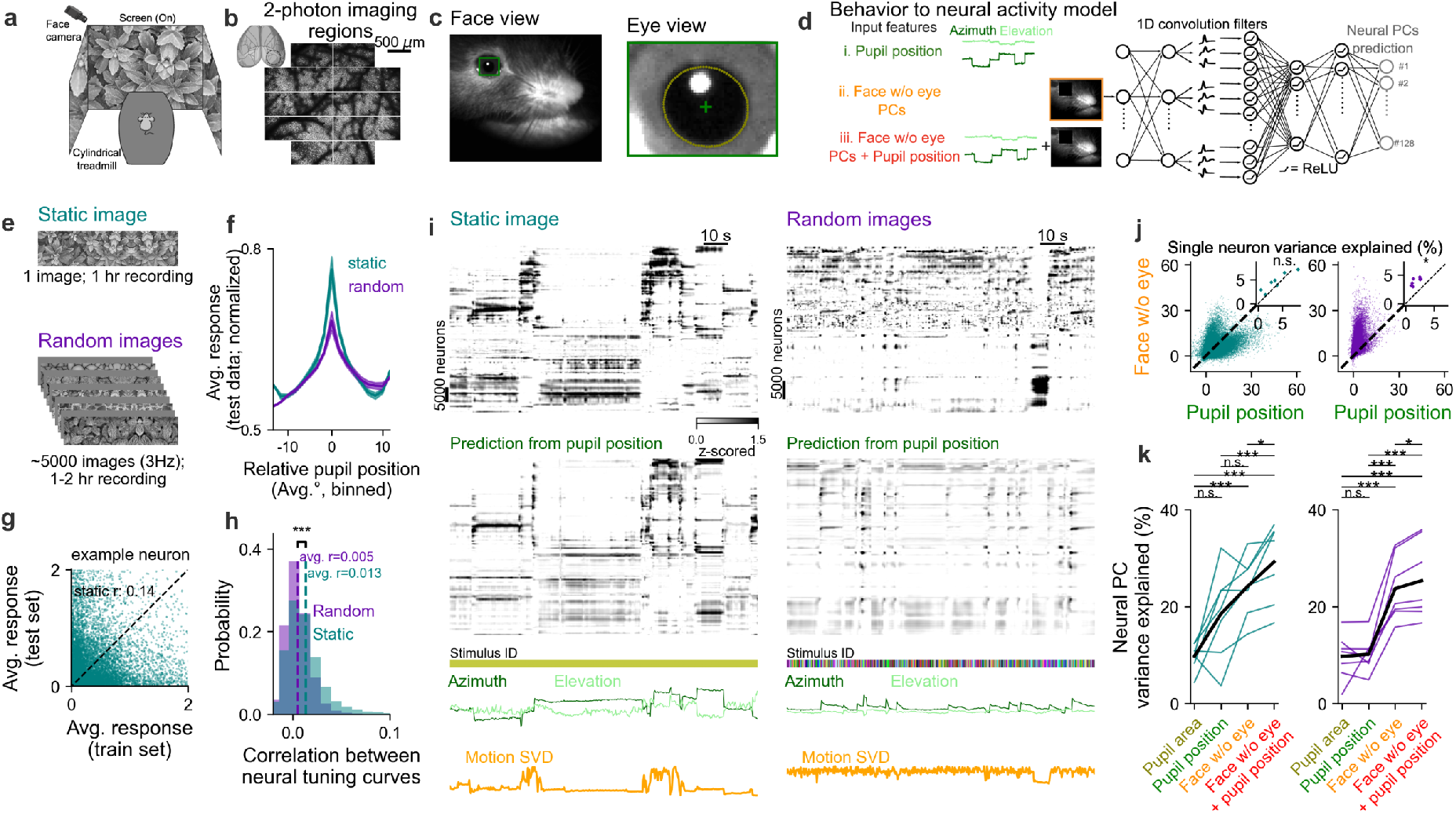
Relative impact of eye and orofacial movements in the presence of retinal input. **a**, Experimental setup. **b**, Neural activity recording in mouse visual cortex using mesoscopic two-photon calcium imaging. **c**, Behavioral video recording of mouse face (left) and zoomed-in view of pupil with ellipse fit (right). **d**, Behavior-to-neural activity prediction using different sets of features. **e**, Image presentation conditions. **f**, Average neural tuning curves after centering on peak response position on training trials. **g**, Example single neuron responses on train vs test trials illustrating *r* calculation. **h**, Histogram of *r* values for random (n=300,317 neurons) and static (n=350,680 neurons) conditions (two-sided t-test; *p* <0.05). **i**, Neural activity Rastermaps for two example recordings in the two conditions. Prediction of neural activity from pupil position shown in the bottom rastermap. Pupil position and first PC of face motion shown for the relevant time periods. **j**, Single-neuron variance explained by model in **d** using Face w/o eye PCs vs. pupil position for example sessions in **i**. Inset shows average variance explained across neurons for each recording (two-sided t-tests; *p*=0.108 (left) and *p*=1.72 × 10^−5^ (right), n=7 mice). **k**, PC variance explained across different models for the two conditions (two-sided t-test, n=7 mice). Colored lines show individual animals and black is average.

To disentangle the effect of eye movements from the effect of visual inputs, we used two visual paradigms: (1) a fixed static image, where eye position and visual inputs were coupled, as in [1]; and (2) random images, where they were uncoupled. For the first condition, we showed the same image for about an hour, while for the second condition we showed ∼5,000 images over an equivalent duration of time (Figure 1e). We started by computing cross-validated tuning curves (see Methods) as a function of pupil position for each neuron (Figure 1f). Since mice primarily make horizontal eye movements when head-fixed [22], we focused on pupil azimuth (Figure S1). The tuning curves for the static image condition had higher peak responses and were more consistent across trials, possibly reflecting direct retinal inputs (example neuron in Figure 1g; all recorded neurons summarized in Figure 1h).

In both conditions, neural population activity underwent global changes at many time points during the recordings (Figure 1i). These were tightly-locked to behavioral state transitions, during which most behavioral readouts underwent changes, including eye position. To quantify how much each behavior contributed to the neural responses, we estimated the variance explained from models using the face and pupil position as inputs. In the static image condition, we found no significant difference in variance explained by face movements (excluding the eye) versus pupil position, at both the single-neuron level (4.5 % vs 3.7 %, Figure 1j, Figure S2) and at the level of population dynamics (24.1% ± 6.2% vs 18.5% ± 9.3% Figure 1k). This result is similar to [1], and is likely due to the tight relationship between retinal inputs and behaviors. When pupil position was decoupled from retinal inputs during the random image session, we found that face movements had better predictions of single-neuron responses compared to pupil movements at the single-neuron (3.9% vs 1.7% Figure 1j, Figure S3) and at the population level (23.7% ± 6.7% vs 10.2% ± 3.7% Figure 1k). Adding pupil position as an additional predictor to the face motion marginally helped prediction in both conditions, and pupil area had similar predictive power to the eye movements. These results were consistent across visual cortical regions (Figure S4). Overall, these experiments show that face movements predict a large portion of neural activity variance independently of eye movements, while eye movements may have predictive power either: 1) indirectly through correlation with retinal inputs; 2) indirectly through correlation with behavioral states; or 3) directly through feedback or corollary discharge modulation.

To further distinguish between these possibilities, we next recorded neural activity in the absence of visual input and performed similar analyses. To track pupil position in complete darkness, we used carbachol eye drops which constrict the pupil allowing identification of its center (Figure 2a). We did not see an effect of using carbachol on the variance of the neural PCs (Figure 2b), qualitatively in the Rastermap plots (Figure 2c), or in the variance explained by the models (Figure 2d). Compared to stimulus conditions (Figure 1), pupil position explained substantially less variance, and much less than face movements (5.4% ± 2.8% vs 21.5% ± 5.1%) (Figure 2d), with no significant increase when using both predictors together (22.1% ± 4.8%). This difference was visually clear in the Rastermap plots, when displaying the predictions from face movements and pupil movements (Figure 2c). Even when predicting the single largest PC, the face prediction model substantially outperformed the model based on pupil position alone (Figure 2e, two-sided paired t-test; p=0.019, n=4 mice), consistent with the hypothesis that pupil-based prediction is primarily due to correlation with the top neural PC which encodes overall arousal levels [7].

**Figure 2:**
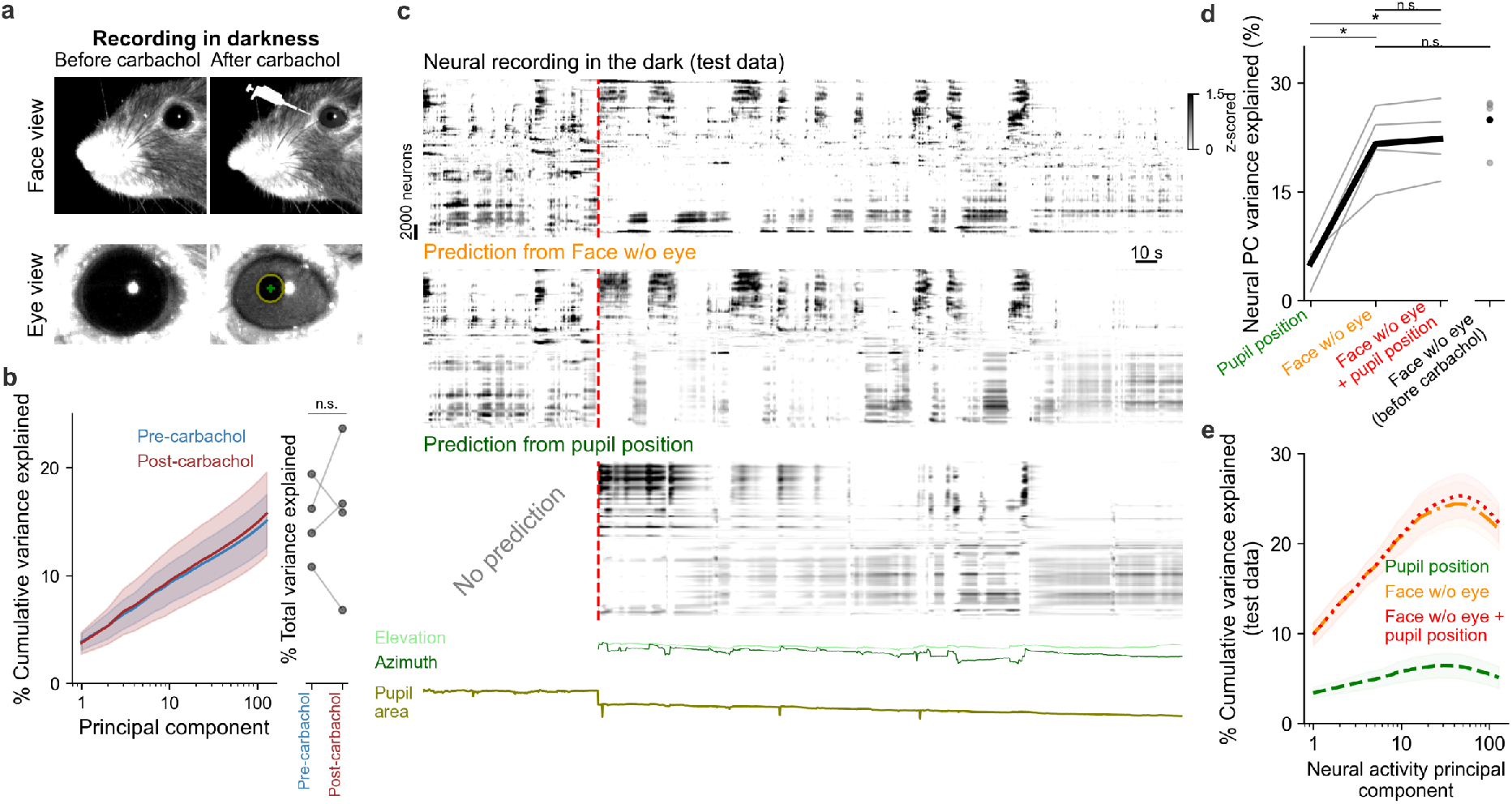
Relative impact of eye and orofacial movements without retinal input. **a**, Face and eye view of mouse before and after carbachol application. **b**, Cumulative (left) and total (right) variance explained by principal components pre- and post-carbachol (left: mean ± s.e.m., n = 4 mice, right: two-sided t-test, p=0.825). **c**, Example rastermap before and after carbachol (top). Prediction of neural activity using Face w/o eye PCs (middle) or pupil position (bottom). Pupil position and pupil area shown below. **d**, Variance explained across PCs by pupil position and face components. Individual animals shown in grey (two-sided t-test; n=4 mice). **e**, Cumulative variance explained across neural PCs (mean ± s.e.m., n = 4 mice).

Previous studies have found that eye movements i.e. saccades are correlated to overall head movements, but did not characterize the relationship between saccades and orofacial movements [22, 23]. To investigate this, we first asked whether orofacial behaviors are coincident with saccades. We performed saccade detection on the pupil position data using a fine-tuned deep learning model [24]. We used keypoints from the Facemap tracker to study orofacial behaviors, such as sniffing and whisking, independently (Figure 3a). We found increased whisker and nose movements around the time of saccades with movements peaking at the saccade times (Figure 3b,c). Pupil area was also increased at these times, and slight running acceleration was observed at saccade times (Figure 3d,e). Conversely, we found that mice performed saccades with a high probability during sniffing (0.078 ± 0.046) and whisking (0.136 ± 0.067) compared to the overall duration of the session (0.009 ± 0.006) (Figure 3f). Similar to previous work [25], we found that saccade direction along the naso-temporal axis was correlated to the direction of whisking and nose movements (Figure 3g). Finally, we used a nonlinear model to predict keypoints from other sets of keypoints (Figure 3h). Whisker and nose position were highly-predictable from each other. Horizontal pupil position was also predictable from whisker and nose position, though at lower levels of variance explained. Overall, we found substantial dependencies between different behavioral variables, making it difficult to determine whether eye position makes a unique, direct contribution to neural activity prediction.

**Figure 3:**
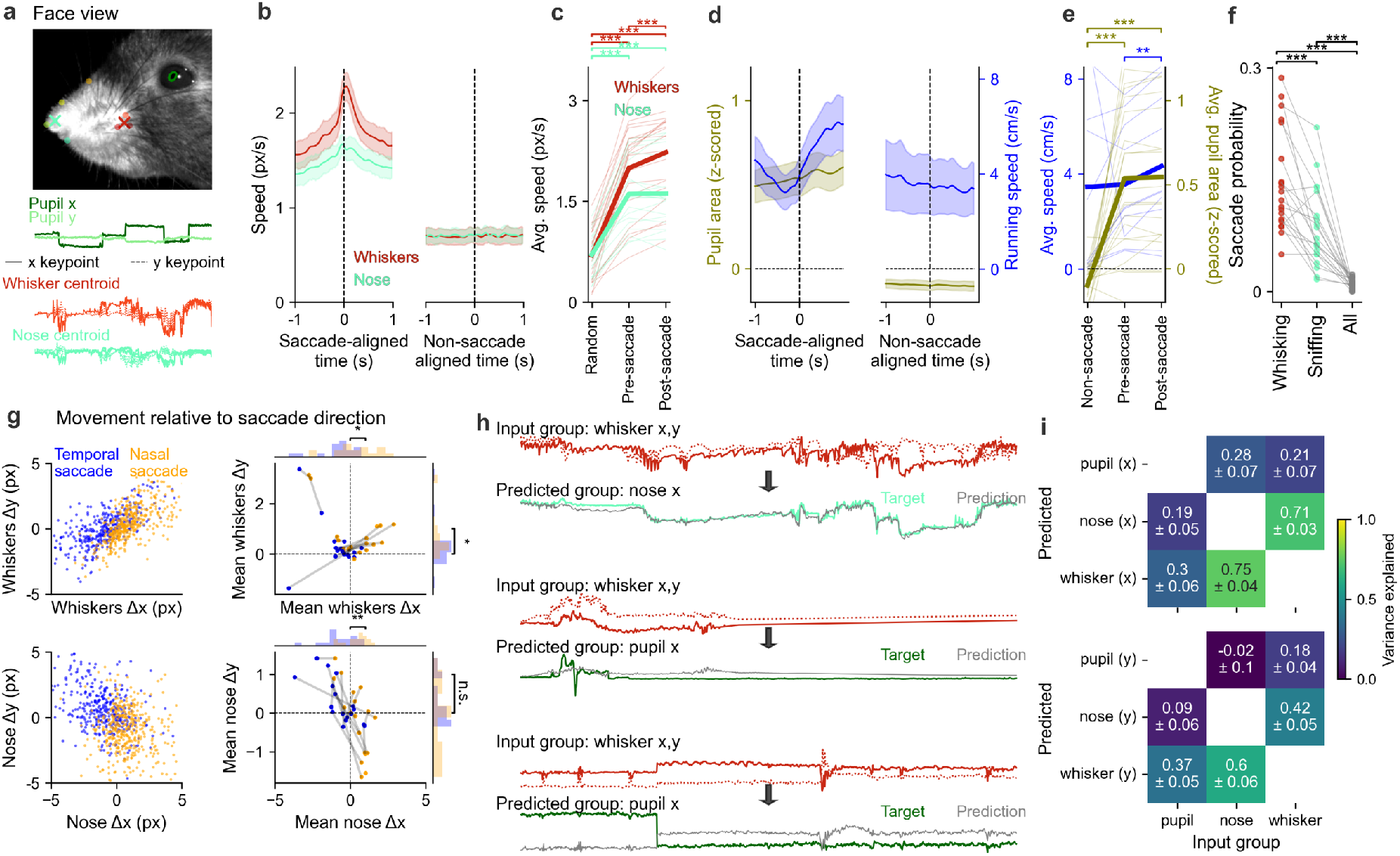
Orofacial movements and pupil dynamics are coupled. **a**, Keypoint tracking of whiskers and nose with pupil annotations in green and centroids (X). Example camera frame (top) and traces (bottom). **b**, Saccade-aligned average whisker speed (left) and aligned to random times (right) (mean ± s.e.m., n=15 sessions). **c**, Average whisker and nose speed during 0.2 s of random / pre-saccade / post-saccade time. Individual animal sessions in thin lines (n=15 sessions, two-sided t-tests between conditions). **de**, Same as **bc** for pupil area and running speed. **f**, Probability of saccade during whisking or sniffing or during all times (two-sided t-tests between conditions). **g**, Whisker and nose velocity relative to saccade direction. Example single-session (left) and average across saccades shown across mice (right, paired t-test). **h**, Example keypoint traces and prediction from other keypoint groups. **i**, Variance explained for x-(top) and y-(bottom) coordinates for different combinations of input and predicted keypoints.

To summarize, we found that high-dimensional orofacial movements are better predictors of responses in the visual cortex of mice, with modest (if any) unique contributions from the eye. Recent studies suggest that orofacial behaviors and their representations might be related to motor intention [26], emotional state [27], or even body state [28]. In addition, prior work has shown that the temporal structure of movement patterns encodes latent cognitive state/variables that are tightly coupled to neural activity [29, 30]. However, the role and origin of these behavioral signals were studied in motor-related areas. In the visual cortex, the discrepancy between the mouse results reported here and primate results [1, 10] may be related to the specific task context in which those datasets were recorded. In [1], monkeys had to fixate to initiate and perform the task, which may require a suppression of motor-related activity across the brain, including in the visual cortex. In contrast, monkeys were locomoting in the [10] study, but face motion signals were not analyzed directly. Locomotion led to a slightly suppressive effect on activity in central regions of V1 [10], but increased responses in peripheral V1 [11], similar to mice, highlighting the need to further investigate extra-foveal visual regions in primates. We also point out that the prediction of neural activity from facial behaviors improves dramatically with the number of simultaneously-recorded neurons [9], and these numbers are relatively low in primate experiments. Thus, it remains possible that spontaneous behaviors could drive neural activity in primate visual cortex during more naturalistic contexts [31] and possibly in darkness, but large-scale recordings will need to be obtained to settle this question definitively.

## Acknowledgments

This research was funded by the Howard Hughes Medical Institute at the Janelia Research Campus. We thank Michalis Michaelos for help with recordings. From the Vivarium, we thank Jim Cox, Crystall Lopez, Anne Kuzspit, Miriam Rose, Alexa Gracias, Gillian Harris, Sarah Lindo, and their respective teams for animal breeding, husbandry, and surgeries. From JeT, we thank Daniel Flickinger, Vasily Goncharov, Alex Sohn, Tobias Goulet, and Steven Sawtelle for help with rig maintenance and upgrades. From MBF Bioscience we thank Georg Jaindl, Mitchell Sandoe, and Boris Djiguemde for scanimage support. We thank the GENIE project team, Caiying Guo and Xianling Zhao at Janelia for generating the TetO-jGCaMP8s transgenic mice.

## Author contributions

A.S., C.S. and M.P. designed the study and wrote the manuscript, with input from all authors. A.S. performed data analysis. M.N and L.Z. performed data collection with help from A.S.

## Data availability

Data will be made available upon publication in a journal.

## Code availability

Code will be made available upon publication in a journal.

## Methods

All experimental procedures were conducted according to IACUC, and received ethical approval from the IACUC board at HHMI Janelia Research Campus. Data analysis and model fitting were performed in python using pytorch, numpy, opencv, scipy, scikit-learn, Rastermap, Facemap, and UnEye [9, 24, 32– 37], and figures were made using matplotlib and jupyter-notebook [38, 39].

### Data acquisition

#### Animals

We performed 18 recordings in 13 mice either bred to express GCaMP6s in excitatory neurons: TRE-GCaMP6s line G6s2 x CamK2a-tTa (available as JAX 024742 and JAX 003010) [40], or GCaMP8s in excitatory neurons: Ptight GCaMP8s x Camk2a-tTA (available as JAX 037717 and JAX 007004) [41]. These mice were male and female, and ranged from 6 to 11 months of age. Mice were housed in reverse light cycle, and were pair-housed with their siblings before and after surgery. Due to the stability of the cranial window surgery, these mice were used for multiple experiments in the lab.

#### Surgical procedures

Surgeries were performed in adult mice (P35–P125) following procedures outlined in [42]. In brief, mice were anesthetized with Isoflurane while a craniotomy was performed. Marcaine (no more than 8 mg/kg) was injected subcutaneously beneath the incision area, and warmed fluids + 5% dextrose and Buprenorphine 0.1 mg/kg (systemic analgesic) were administered subcutaneously along with Dexamethasone 2 mg/kg via intramuscular route. Measurements were taken to determine bregma-lambda distance and location of a 4 mm circular window over V1 Cortex, as far lateral and caudal as possible without compromising the stability of the implant. A 4+5 mm double window was placed into the craniotomy so that the 4mm window replaced the previously removed bone piece and the 5mm window lay over the edge of the bone. After surgery, Ketoprofen 5mg/kg was administered subcutaneously and the animal allowed to recover on heat. The mice were monitored for pain or distress and Ketoprofen 5mg/kg was administered for 2 days following surgery.

#### Videography

The camera setup was similar to the setup in [9]. The entire setup was enclosed in a large black box to prevent light from the room from entering the microscopy light path and from entering the mouse’s eye (measured as fully dark - 0.00 lux, see [9] for details). A Thorlabs M850L3 - 850 nm infrared LED was pointed at the face of the mouse to enable infrared video acquisition in darkness. The wavelength of 850nm was chosen to avoid the 970nm wavelength of the 2-photon laser, while remaining outside the visual detection range of the mice [43, 44]. The videos were acquired at 35 or 50 Hz using a FLIR camera with a zoom lens and an infrared filter (850nm, 50nm cutoff) pointed at the side of the face contralateral to the recording area. The camera acquisition software was a modified version of the Facemap software.

#### Carbachol eye drops for pupil constriction

To track eye movements in the dark, we used a pharmacological approach to constrict the pupil using carbachol eye drops. The eye drops were administered to the eye facing the camera and contralateral to the recording region. Carbachol was diluted in saline solution at 100 mM concentration and subsequently passed through a micron filter to obtain a sterile carbachol solution. Carbachol solution was applied to the eye using a micro-pipette to administer a single drop ranging from 5-15 *µ*L. The effect of the eye drops lasted around 20-30 minutes and in two mice another 5-15 *µ*L drop was administered to obtain a longer neural recording session. It is important to note that carbachol can have other side effects when ingested by the animal. Hence, the protocol was optimized to ensure the drops only reach the eye to avoid any side-effects relating to the ingestion of carbachol. It is advised to start with one drop ranging from 5-15 *µ*L and wait a around 5 minutes to see its effect before proceeding with another drop of 5-15 *µ*L.

#### Imaging acquisition

We used a custom-built 2-photon mesoscope [45] to record neural activity, and ScanImage [46] for data acquisition. We used a custom online Z-correction module (now in ScanImage), to correct for Z and XY drift online during the recording. As described in [42], we used an upgrade of the mesoscope that allowed us to approximately double the number of recorded neurons using temporal multiplexing [47]. The mice were head-fixed and free to run on a cylindrical treadmill. The field of view for recording was selected such that large numbers of neurons could be observed, with clear calcium transients. The mesoscope scanning of each region occurred at 3 Hz sampling frequency.

#### Processing of calcium imaging data

Calcium imaging data was processed using the Suite2p toolbox [48], available at www.github.com/MouseLand/suite2p. Suite2p performs motion correction, ROI detection, cell classification, neuropil correction, and spike deconvolution as described elsewhere [7]. Spike deconvolution was carried out using a new algorithm implemented in Suite2p [48], which employs a neural network trained to predict ground-truth spikes from calcium traces [49]. We obtained 42,476 ± 14,020 (s.d., n=18 session recordings in 13 mice) neurons in the recordings.

#### Visual stimulus

Neural recordings were conducted in three different visual stimulus conditions: (i) presentation of a static image, (ii) presentation of random images, (iii) and in darkness. Visual stimuli were presented on three monitors shown in Figure 1a. In (i) a single static image was kept on the screens throughout the duration of the neural recording (Figure 1e). In (ii) random images were presented for 139 ms each followed by 139 ms of gray screen, resulting in a presentation rate of 3.6 Hz (Figure 1e). In (iii) recordings were done in the dark (without visual input) with the monitors turned off.

### Behavior tracking

#### Acquisition of face videos

Behavioral data including face videos were collected at a sampling frequency of ∼ 35 Hz or ∼ 50 Hz. We saved the full face video downsampled by a factor of 2, with a resolution of 320 by 400 pixels, while the cropped eye region video was saved at full resolution (around 70 by 45 pixels). (Figure 1c).

#### Eye tracking

The Facemap software package was used to track pupil position [9]. The graphical user interface (GUI) of Facemap was used to load the eye video and to draw an elliptical region of interest (ROI) around the inner area of the eye which contains the pupil. The ROI computes an ellipse around the pupil area based on the contrast/saturation difference with the rest of the eye. A multivariate Gaussian is fitted around the pixels in the box using maximum likelihood, and the pupil position (x,y) is defined as the mean of the Gaussian. The pupil area is defined as the area of an ellipse at 2 standard deviations away from the mean of the Gaussian. A pupil area trace is obtained from processing the entire video (Figure 1c). We removed outliers from the pupil area and x,y traces, by performing median-filtering with a window of 30 frames on the traces and replacing outliers with the median-filtered values. Outliers were defined at timepoints where the difference between the raw trace and the median filtered trace was greater than half the standard deviation of the raw trace.

The polar coordinates of the pupil position (azimuth and elevation) were obtained to get directionality of eye movement (Figure 1i). The polar coordinates were calculated using a user-defined axis extending from the left and right eye keypoints of the mouse (Figure S1b). The position of the pupil was used to compute its displacement relative to the user-defined eye axis (Figure S1c). Saccade directions, i.e. nasal versus temporal, were defined according to the polar coordinates (Figure S1f).

#### Saccade detection

Saccade detection was performed using the u’n’eye model [24]. Inputs to the model were pupil x,y coordinates sampled at 35-50 Hz from the Facemap software [9]. The pupil position trace was split into segments of length 1000 frames with padding of 15 frames around each sequence. Data was randomly divided into train and test sets using a 80/20 split. Next, a base model was loaded with pre-trained ‘Andersson weights’ to obtain saccade predictions for two light recordings. The predicted saccade labels were refined (post-hoc) using software in Matlab [50]. The refined labels were subsequently used to fine-tune the base model, ‘Andersson weights’, using learning rate 5*e* − 4 and default hyperparameters. Finally, saccade labels for remaining light recordings were obtained using predictions of the fine-tuned model and post-hoc curation was performed for each session in the Matlab software [50]. Saccade times obtained in behavioral timescale were plotted on pupil position traces and resampled to neural timescale for plotting on neural data (Figure 1i, Figure S1a).

In the dark recordings, this fine-tuned model did not perform well, due to the changing baseline from the carbachol. Therefore, we manually labeled the saccades in the dark recordings using an updated version of the Facemap GUI.

Across recordings, a linear relationship was observed for saccade amplitude versus saccade velocity, suggesting accurate saccade time prediction (Figure S1e).

#### PCA of behavioral videos

Singular value decomposition of the mouse face videos was performed using Facemap, as described previously [9]. For each session, we manually defined an ROI surrounding the mouse face, and another ROI to exclude pixels covering the eye, resulting in the ‘face w/o eye’ PCs (Figure 1d). We used 500 movie PCs for each video, and each of these PCs was scaled by its singular value.

### Tuning curves

Tuning curves for the light recordings were computed to evaluate neural response to visual stimuli (Figure 1f). The tuning curve was computed using pupil position (azimuth) because mice made most saccades in the nasal-temporal direction (Figure S1f). The pupil azimuth values were resampled to neural timestamps and then binned into nine percentile-based bins so each bin contained an equal number of samples. Subsequently, each bin of pupil position was divided into a 50-50 split of random train and test timepoints. The activity of each neuron was normalized by its standard deviation across all timepoints and plotted during train and test trials (Figure 1g). We computed the correlation between train and test timepoints across all bins as an estimate of the reliability of the responses of each neuron (Figure 1gh). Preferred pupil position bin for each neuron was defined as the maximum average response across bins using only training timepoints per bin. The average responses on test trials, i.e. the tuning curves, were aligned across neurons using the preferred stimulus bin of each neuron from the average responses on train trials (Figure 1f).

### Behavior to neural prediction

#### Neural data

Deconvolved neural activity traces were z-scored: the activity of each neuron mean-subtracted and divided by its standard deviation. PCA of neural data was performed to obtain 128 PCs for each recording, and each PC was scaled by its singular value to retain the variance distribution in the original data. The PCs were used to reduce the neural data dimensionality and improve SNR for fitting a model from behavior to neural activity.

The neural and behavioral recordings consisted of multiple blocks for one session that were concatenated for analysis. The light recordings consisted of a single block in each session. The dark recordings consisted of two or three blocks in 2 mice for each session: before carbachol, after carbachol and (optionally) another after carbachol block. The length of recording can affect the performance of the model, so similar length of time was used for light recordings by cropping all sessions to the same length (∼ 1h 15 mins) and ∼ 1h recordings collected in the dark.

#### Model architecture

Behavior to neural activity prediction was performed using the Facemap deep neural network model [9]. In brief, the model consists of a linear layer, a 1D temporal convolution with relu, two fully-connected layers with relu’s, and a linear projection to the output layer (Figure 1d). The three types of inputs to the model were: (i) pupil position: azimuth and elevation (2 features), (ii) Face w/o eye PCs (500 features), (iii) Face w/o eye PCs + pupil position (502 features). The outputs of the model were the 128 PCs of neural data. The first linear layer output was set to a size of 5 for pupil position and pupil area predictions (because the input features were less than 10), and set to a size of 250 for other behavioral predictions using input features>10. For temporal convolution, default filter size of 201 timepoints was used (Figure 1d). A separate model was trained on each recording session. For light recordings, one long single block was used to train the model. For dark recordings, a model was trained across all blocks for face w/o eye prediction, whereas only the after carbachol block(s) were used for pupil position and face w/o eye + pupil position predictions.

#### Model training

The model was trained using k-fold (k=5) cross-validation. Each of the splits consisted of the behavioral data divided into continuous time segments of length ∼ 67s, with ∼ 2-2.8 seconds of padding between segments. Each of these segments were assigned to the training, validation and test sets, with probabilities of 60%, 20% and 20% respectively. Similarly, neural data was divided into the training, validation and test sets using the nearest neural data timestamps to the camera timestamps in each set.

We trained the model with the AdamW optimizer [51] with a learning rate of 1e-3, weight decay of 1e-4, and a batch size of 1. During training, the variance explained (Equation 1) was computed for the validation set. The model was trained for a maximum of 300 epochs and a minimum of 50 epochs using early stopping when validation accuracy, i.e. variance explained, did not improve for 15 epochs. A learning rate scheduler was used to reduce learning rate by a factor of 0.1 when the maximum accuracy on validation set did not improve for 10 epochs, before 50 epochs were reached. The model with the lowest validation accuracy was saved and used for test set inference.

#### Model evaluation

The model prediction accuracy was evaluated on the test set using the variance explained across the neural PCs (VE_PC_), defined as

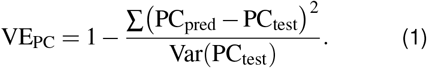

Cumulative variance explained for each neural PC (CVE) was computed by weighting the predicted variance explained for each neural PC by the fraction of total neural variance captured by that PC (normalized variance):

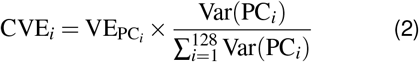

We computed the single-neuron variance explained by multiplying the predicted PC activity traces by the neural PC vectors, as in [9].

### Orofacial behaviors and eye movement

The Facemap tracking network was used to track orofacial keypoints on the mouse face (Figure 3a). Keypoint predictions were obtained using the default base Facemap network, then 10 video frame predictions were corrected and used to train a fine-tuned model for each mouse as described in [9].

Keypoint predictions for different bodyparts in each group (whiskers and nose) were averaged to obtain a centroid position for the group denoted as X in Figure 3a. For each bodypart centroid speed was calculated as the Euclidean distance between consecutive frames. Whisker and nose centroid speeds were then aligned to the starts of saccades (Figure 3b). The running speed of mouse and pupil area (z-scored) were also aligned to the starts of saccades (Figure 3d). To average across recordings (which varied in sampling rate from 35-50Hz), we resampled the behavioral variables and saccade times to 50 Hz. As a control for comparison with saccade time periods, random continuous time segments of the same length excluding saccades were used to align and average pupil area, running speed, whisker and nose centroid speed (Figure 3bd right). A 0.2 s segment of time was selected before the start of saccade time (pre-saccade), 0.2 s after start of saccade time (post-saccade), and 0.2 s of non-saccade aligned time. Finally, the average change in centroid speed, pupil area or running speed was compared during the non-saccade, pre-saccade and post-saccade times for each mouse (Figure 3ce).

Saccade probability during different behaviors was computed using keypoints data. A centroid speed above 97^*th*^ percentile was used to define whisking and sniffing times. To obtain saccade probability, the number of time points covered by saccade region (start and end of saccade) was summed and divided by the total number of time points for whisking, sniffing, and all timepoints (Figure 3f). This computed how likely the mouse was to perform a saccade when the animal is whisking, sniffing, versus during the entire session.

Movements of orofacial keypoints were obtained relative to saccade direction. Saccade direction was defined as nasal versus temporal according to the direction of change of the eye azimuth. Next, the centroid positions at the start and end of saccade were used to compute two values: change in x-position (Δ*x*) and change in y-position (Δ*y*) for each mouse session (Figure 3g left). Average centroid movements during the two saccade directions for all mice were used to perform a paired t-test (Figure 3g right).

#### Behavior to behavior prediction model

We developed a temporal convolutional network to perform prediction of one behavior centroid (whisker, nose or pupil) from another (Figure 3h). The network architecture was designed to capture behavioral dependency at different timescales using varying receptive field sizes. The architecture consisted of an initial projection layer, a stack of dilated residual blocks with grouped convolutions, and a final linear projection to the output space. Input to the model was either pupil *x, y* position or keypoint centroid *x, y* position, and the output was pupil *x, y* position or keypoint centroid *x, y* position.

The input to the model was first passed through a convolutional layer (projection layer) consisting of 32 channels with kernel size 1. To capture behaviors at different timescales, we used a stack of three residual blocks with a channel size of 32 and with increasing dilation factors (*d*=1,2,4), similar to previous work [52, 53]. Each residual block consisted of: (i) a 1D convolution with kernel size of 51 (representing 1 or 1.4s of time for data collected at 35 or 50 Hz respectively) and dilation *d*, (ii) batch normalization and ReLU activation, (iii) dropout (rate = 0.2) for regularization, (iv) and a convolutional layer with kernel size 1. The output of the block was then added to the input of the block before passing through another ReLU activation. The convolution in the first residual block was initialized with Gabor wavelets at various frequencies. Finally, the output from the sequence of residual blocks was passed through a 1 × 1 convolution to project the channels to the output dimension of 2.

#### Model training and evaluation

The inputs and outputs were partitioned into contiguous segments of length 2000 and padding of 50 frames (1-1.4s). Train and test data segments were then randomly selected using an 80/20 split. The loss function was the mean-squared error between the predicted and ground-truth segment. The model parameters were optimized using backpropagation with the AdamW optimizer with learning rate 1*e* − 3 and and weight decay of 1*e* − 2 for 200 epochs with a batch size of 4. A mean-squared error (MSE) loss function was computed on each epoch for training data as follows:

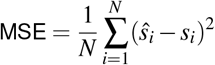

Where *ŝ*_*i*_ and *s*_*i*_ denote the predicted segment and ground truth segment for sample *i*, respectively.

To quantify model performance, we computed variance explained between predicted and true sequences. The variance explained for each output channel (x, y coordinate) was computed as:

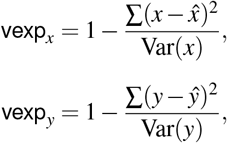

where 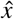 and ŷ denote model prediction and *x, y* are the ground truth. For each test segment, predictions corresponding to the first and last 50 time points (frames) were excluded from evaluation to minimize the effect of padding artifacts. The variance explained was averaged across all sessions for each of the input predictors (Figure 3i).

---

**Supplementary data 1:** Pose tracking and saccade detection video.

**S1:**
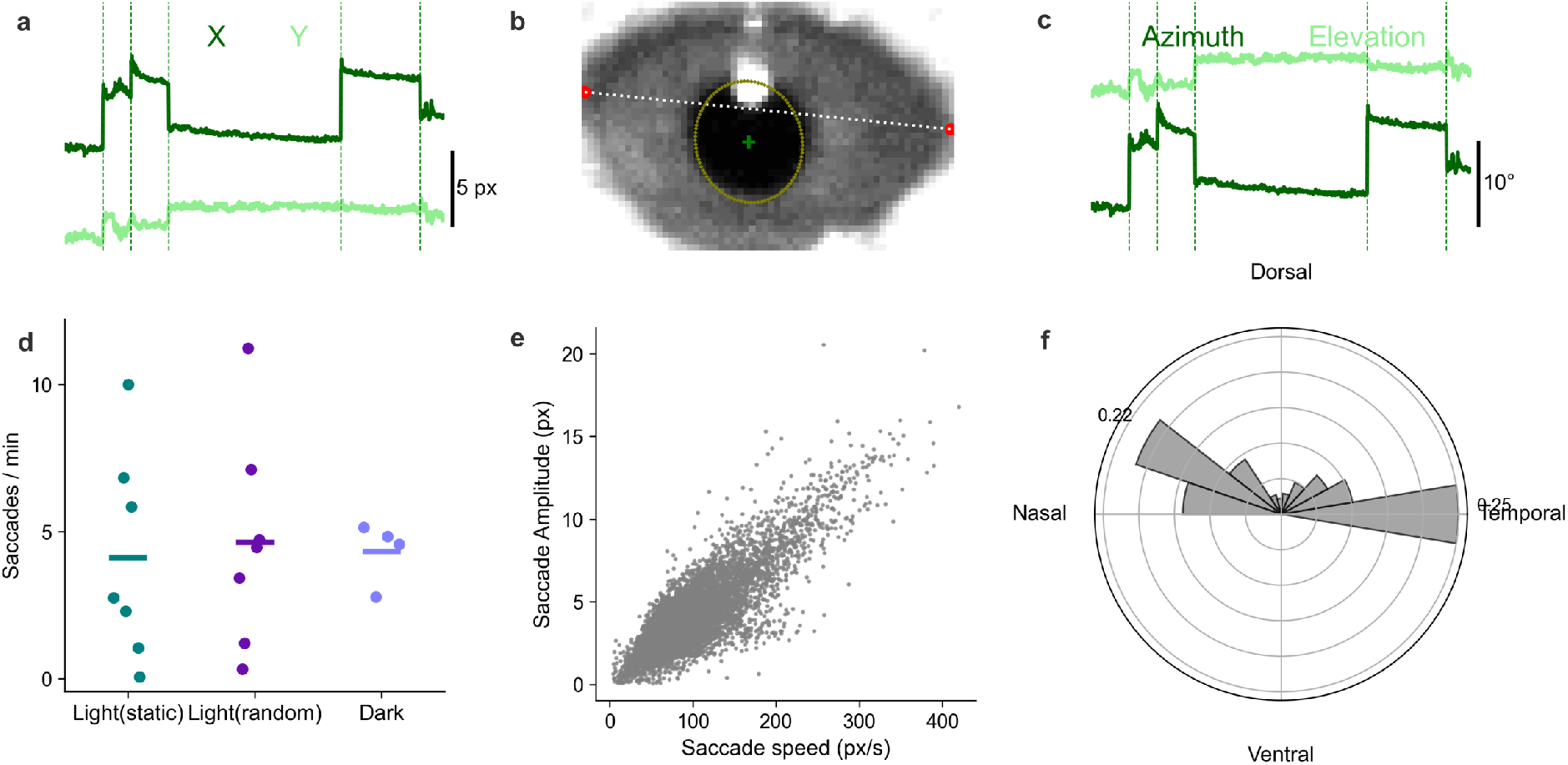
Polar coordinate transformation of pupil x,y coordinates and saccade statistics. **a**, Pupil position x,y coordinates of example video. Green vertical dotted lines indicate saccade time. **b**, Eye view of example session. Ellipse in olive highlights pupil area and green + shows pupil center of mass used to obtain x,y coordinates. Red dots show user-defined eye keypoints used to obtain eye axis along the nasal-temporal direction. **c**, Transformation of pupil x,y coordinates to polar coordinates showing angles in degrees for azimuth and elevation. **d**, Saccade frequency (saccades/minute) for light static (n=7 mice), light random (n=7 mice), and dark (n=4 mice). Horizontal line shows mean across mice. **e**, Saccade amplitude (px) versus saccade speed (px/s) (defined as maximum speed during saccade). Each scatter point represents a saccade; data shown for all sessions. **f**, Probability of saccades recorded in nasal, dorsal, temporal or ventral direction determined from polar coordinates.

**S2:**
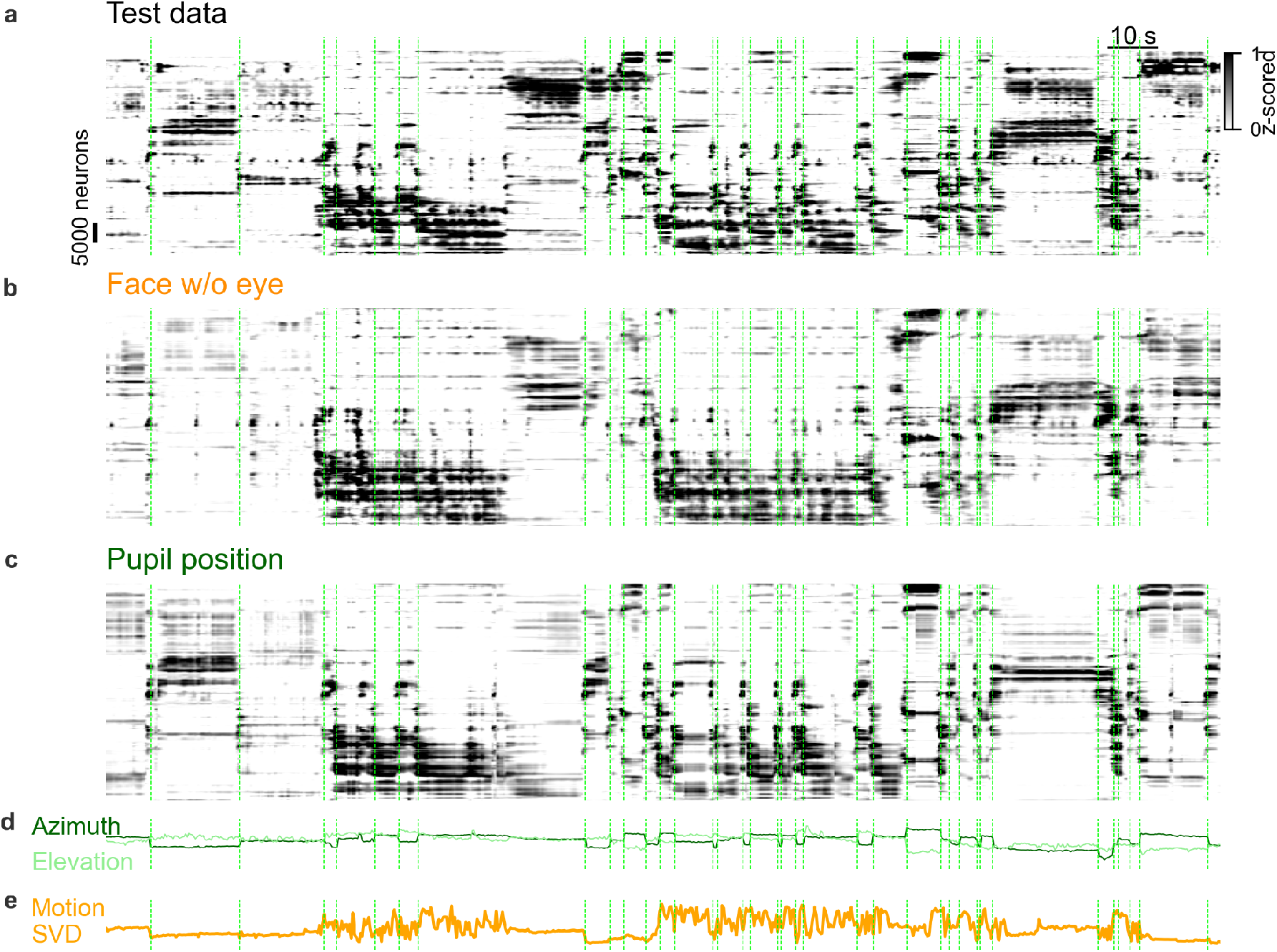
Neural activity prediction during static image presentation using Face w/o eye PCs and pupil position. **a**, Rastermap sorting of neural activity recorded in the visual cortex of example mouse during static stimulus presentation. Green vertical dotted lines indicate saccade time. **b**, Prediction of neural activity using Face w/o eye PCs. **c**, Prediction of neural activity using pupil position. **d**, Pupil elevation and azimuth traces shown in green. **e**, First PC of behavioral video of mouse face.

**S3:**
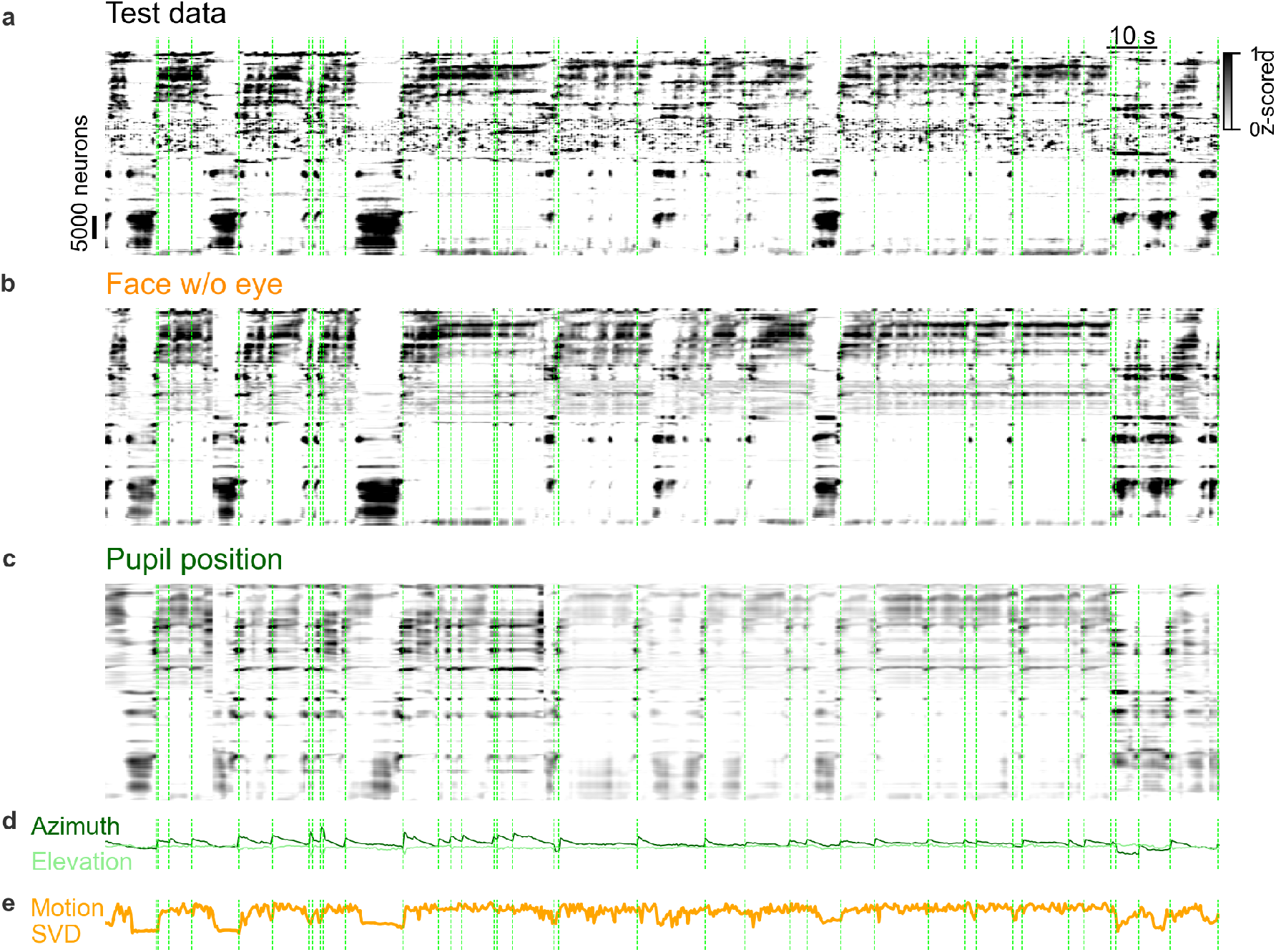
Neural activity prediction during random stimuli presentation using Face w/o eye PCs and pupil position. **a**, Rastermap sorting of neural activity recorded in the visual cortex of example mouse during random stimuli presentation. Green vertical dotted lines indicate saccade time. **b**, Prediction of neural activity using Face w/o eye PCs. **c**, Prediction of neural activity using pupil position. **d**, Pupil elevation and azimuth traces shown in green. **e**, First PC of behavioral video of mouse face.

**S4:**
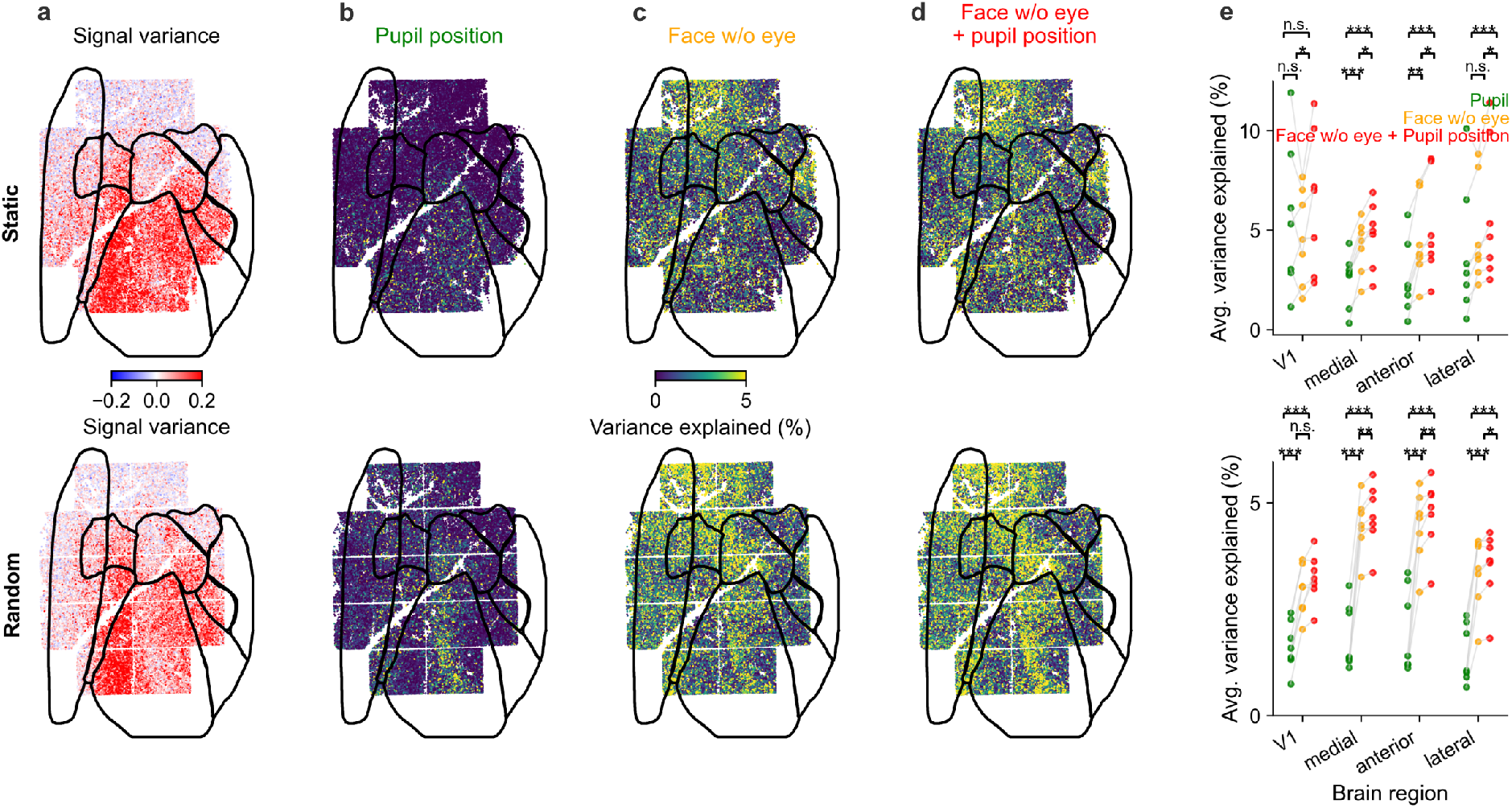
Signal variance and single-neuron variance explained by Face w/o eye PCs and pupil position across different regions of visual cortex during static stimulus and random stimuli presentation. **a**, Signal variance of single neurons in V1, medial, anterior and lateral regions of visual cortex during static [top] and random [bottom] stimuli presentation. **b**, Single-neuron variance explained by pupil position during static [top] and random [bottom] stimuli presentation for an example mouse. **c**, Single-neuron variance explained by Face w/o eye PCs during static [top] and random [bottom] stimuli presentation. **d**, Single-neuron variance explained by Face w/o eye PCs + pupil position during static [top] and random [bottom] stimuli presentation. **e**, Average variance explained across neurons by behavioral variables in V1, medial, anterior and lateral regions of mouse visual cortex shown for static [top] and random stimuli [bottom] recordings.

